# Pervasive generation of non-canonical subgenomic RNAs by SARS-CoV-2

**DOI:** 10.1101/2020.04.28.066951

**Authors:** Jason Nomburg, Matthew Meyerson, James A. DeCaprio

## Abstract

**Background:** SARS-CoV-2, a positive-sense RNA virus in the family *Coronaviridae*, has caused a worldwide pandemic of coronavirus disease 2019 or COVID-19 Coronaviruses generate a tiered series of subgenomic RNAs (sgRNAs) through a process involving homology between transcriptional regulatory sequences (TRS) located after the leader sequence in the 5’ UTR (the TRS-L) and TRS’ located near the start of structural and accessory proteins (TRS-B) near the 3’ end of the genome. In addition to the canonical sgRNAs generated by SARS-CoV-2, non-canonical sgRNAs (nc-sgRNAs) have been reported. However, the consistency of these nc-sgRNAs across viral isolates and infection conditions is unknown. The comprehensive definition of SARS-CoV-2 RNA products is a key step in understanding SARS-CoV-2 pathogenesis.

**Methods:** Here, we report an integrative analysis of eight independent SARS-CoV-2 transcriptomes generated using three sequencing strategies, five host systems, and seven viral isolates. Read-mapping to the SARS-CoV-2 genome was used to determine the 5’ and 3’ coordinates of all identified junctions in viral RNAs identified in these samples.

**Results:** Using junctional abundances, we show nc-sgRNAs make up as much as 33% of total sgRNAs *in vitro*, are largely consistent in abundance across independent transcriptomes, and increase in abundance over time during *in vitro* infection. By assessing the homology between sequences flanking the 5’ and 3’ junction points, we show that nc-sgRNAs are not associated with TRS-like homology. By incorporating read coverage information, we find strong evidence for subgenomic RNAs that contain only 5’ regions of ORF1a. Finally, we show that non-canonical junctions change the landscape of viral open reading frames.

**Conclusions:** We identify canonical and non-canonical junctions in SARS-CoV-2 sgRNAs and show that these RNA products are consistently generated across many independent viral isolates and sequencing approaches. These analyses highlight the diverse transcriptional activity of SARS-CoV-2 and offer important insights into SARS-CoV-2 biology.

## Background

Severe acute respiratory syndrome coronavirus 2 (SARS-CoV-2) emerged from Wuhan, China in December 2019 and rapidly led to a world-wide pandemic, known as COVID-19 (1). The initial sequencing report found that this virus maintains 89.1% nucleotide identity to SARS-like coronaviruses found in bats in China (1).

Coronaviruses (CoV) including SARS-CoV-2 contain large, ~ 30kb, positive-sense, single-stranded RNA genomes with a unique genome organization structure. The 5’ end of the genome contains two large open reading frames (ORFs), ORF1a and ORF1b. ORF1a produces a large polyprotein, while ribosomal slippage at an RNA pseudoknot structure and slippery sequence at the end of ORF1a occasionally leads to a frameshift and subsequent translation of a joint ORF1a and ORF1b polyprotein (2). ORF1a and ORF1ab polyproteins contain protease activity that can cleave the polyproteins into a variety of nonstructural proteins (Nsp) capable of facilitating viral transcription and replication and modulating host transcription and translation. Following ORF1b, there is an arrangement of ORFs encoding structural and accessory proteins that tend to vary in content and order across the CoV family. While structural proteins are incorporated into emergent virions, accessory proteins are thought to be dispensable for replication *in vitro* but to increase fitness *in vivo* (3–5).

Initial translation of ORF1ab can occur directly from incoming viral genomic RNA, generating the ORF1a and ORF1ab polyproteins and initiating infection. However, because the viral genome is so large, and because downstream ORFs are far from the 5’ end of the genome, effective translation of these ORFs requires the generation of subgenomic RNAs (sgRNAs) (6). To generate these sgRNAs, it is thought that CoV undergoes a process of discontinuous extension during synthesis of negative-stranded RNA templates. As the virus-encoded RNA-dependent RNA polymerase progresses from the 3’ end towards the 5’ end of the viral genome, it encounters transcriptional regulatory sequences (TRS) upstream of each major ORF (TRS-B). Here, the polymerase can skip to a similar TRS that is 3’ of a shared leader sequence within the 5’ UTR of the CoV genome (TRS-L). These antisense sgRNAs are then transcribed, resulting in a series of tiered sgRNAs containing the same 5’ leader sequence (6).

Several studies have characterized SARS-CoV-2 sgRNAs using Nanopore direct RNA sequencing (dRNAseq). In contrast to short-read sequencing, which can produce read pairs of 150bp each, dRNAseq is capable of directly sequencing entire transcripts including the full length of the viral genome (29903 bases). This allows dRNAseq to identify RNA isoforms and variants that may be challenging to deconvolute using short-read sequencing. Using nanopore sequencing, Taiaroa et al. identified eight major sgRNAs, as well as a small number of non-canonical junctions between the 5’ leader sequence and downstream regions of the SARS-CoV-2 genome (7). Kim et al. identified a total of 9 distinct subgenomic RNAs, as well as a series of non-canonical subgenomic RNAs (nc-sgRNAs) that contain unexpected junctions between 5’ and 3’ sequences (8). Using short-read DNA nanoball sequencing data, they confirmed the existence of noncanonical junctions at various sites across the genome (8). Davidson et al. identified a series of transcripts encoding the nucleocapsid (N) protein, but with distinct small internal deletions, and validated this finding with proteomic evidence (9).

While these previous studies identified non-canonical RNA junctions and unexpected RNA species in isolation, there were no integrative analyses of the SARS-CoV-2 junctions. It is unclear if identified non-canonical junctions are broadly representative of SARS-CoV-2 biology or if they are a result of dataset- or isolate-specific artifacts. Here, we report an integrative analysis of eight independent SARS-CoV-2 transcriptomes generated using three sequencing strategies, five host systems, and seven viral isolates to comprehensively characterize the landscape of canonical and non-canonical sgRNAs generated by SARS-CoV-2.

## Results

### Analysis of three independent dRNAseq datasets reveals canonical and noncanonical junctions

To determine the major RNA species present in SARS-CoV-2-infected cells, we applied a uniform computational approach to three independent dRNAseq datasets from Taiaroa et al., Kim et al., and Davidson et al. (7, 10, 11). To assess sgRNA presence, we identified junction-spanning reads following read-mapping against the SARS-CoV-2 reference genome.

Analysis of the 5’ and 3’ locations of junctions separated by at least 1000 bases revealed that the majority of junctions consist of a 5’ end originating in the 5’ UTR and a 3’ end at distinct sites in the 3’ region of the genome (Figure 1A–C). These represent the canonical junctions between the leader sequence and 3’ sequences that generate the major sgRNAs (6). Plotting the counts of 3’ junctions that contain a 5’ end within the first 100 bases revealed eight prevalent junction points along the 3’ end of the genome (Figure 1D–F), all of which are proximal to the TRS core sequence ACGAAC. Interestingly, we observed that the 3’ junction peak and TRS-B closest to ORF6 lands 151 bases within the M ORF. Based on these junction analyses and the locations of identified TRS body sites, predicted sgRNAs and their most 5’ gene or genes are shown in Figure 1G. We predict 8 major sgRNAs based both on identification of the TRS core sequence and presence of a dominant junction peak. In contrast to Kim et al. (8) but in concordance with Taiaroa et al. (7) and Davidson et al. (9), we do not find evidence of a prominent 3’ junction point near ORF7B (Figure 1D–F).

**Figure 1.**
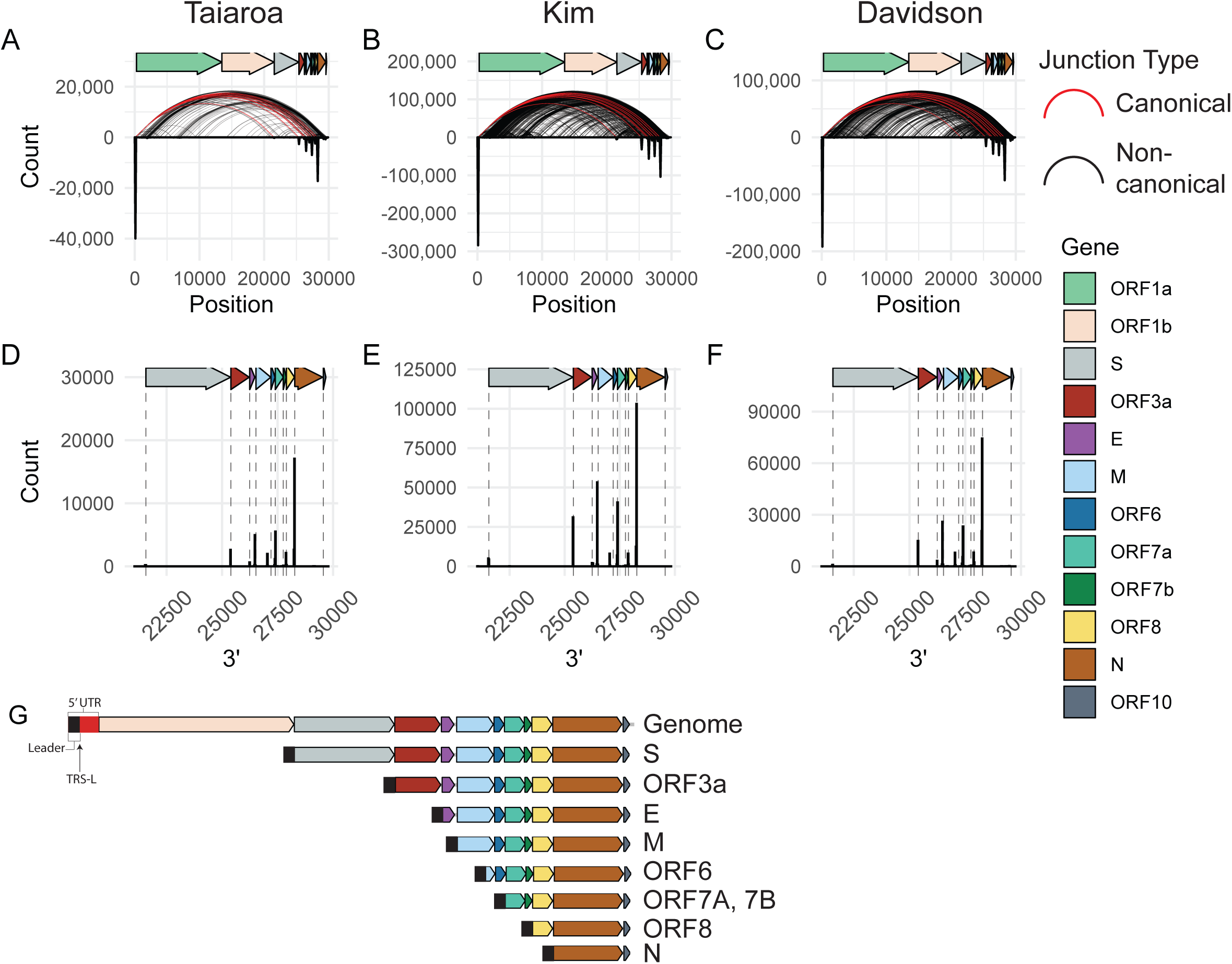
SARS-CoV-2 generates a defined population of canonical subgenomic RNAs. **A-C)** For each location on the viral genome, a histogram of 5’ and 3’ junctions at that position was calculated and plotted as an inverse peak. The histogram bin size is 100 bases, meaning each inverse peak represents the cumulative count of 5’ or 3’ junctions occurring within that span. Curved lines represent the 5’ and 3’ locations of junctions that occur at least twice. Red curves represent canonical junctions, black curves represent non-canonical junctions. “Taiaroa” (7), “Kim” (8), and “Davidson” (9) represent the three independent SARS-CoV-2 dRNAseq datasets investigated. **D-F)** A histogram of 3’ junctions past position 21000 with a 5’ end before position 100 are plotted. Dashed lines indicate the start coordinates of annotated viral genes. Bin-size is 20 bases. **G)** Based on junction analysis, the predicted major species of virus-produced RNAs are represented. The most 5’ gene or genes on each subgenomic RNA is listed.

### Many SARS-CoV-2 junctions are non-canonical

Similar to observations by Kim et al. (8), we identified an unexpectedly diffuse pattern of noncanonical junctions across the genome (Figure 1A–C, black arcs), suggesting that there are many unexpected RNA species. Had there been only canonical junctions, the arcs on this plot would be exclusively between the first 100 bases and defined points on the x-axis (Figure 1A–F).

Because the transcriptomes in these long-read sequencing studies (7, 8, 11) all consist of dRNAseq from infected Vero cells, we sought to confirm this observation in independent transcriptomes from other SARS-CoV-2 isolates, in other host systems, and using other sequencing technologies. To this end, we assessed five additional transcriptomes generated from Vero cells (12), A549 cells (13), Calu3 cells (14), bronchial organoids (15), and ferret nasal washings (13) using Illumina PolyA or total RNA sequencing. Detailed information regarding each sample is detailed in the methods. Junction analysis in these transcriptomes likewise revealed pervasive non¬canonical junctions (Supplementary Figure 1A–E). Quantification of the relative abundance of canonical and non-canonical junctions in these transcriptomes revealed that up to 33% of junctions are non-canonical *in vitro* (Figure 2A, Supplementary Figure 1F). The transcriptome from a ferret’s nasal washing contained 51% non¬canonical junctions, indicating that non-canonical junctions are also produced *in vivo.*

**Figure 2.**
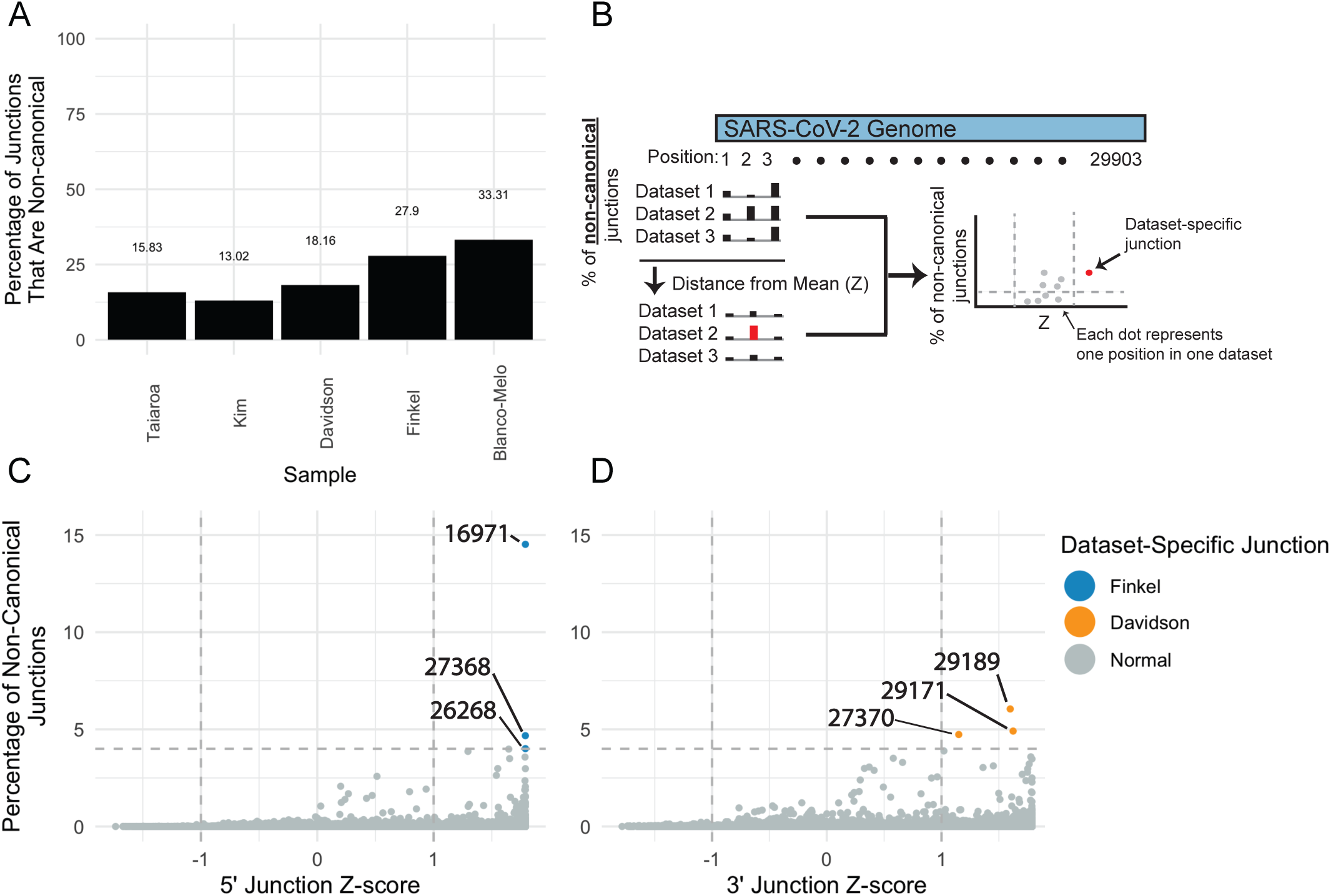
Non-canonical junctions are consistent across independent datasets. **A)** Percentage of junctions that are non-canonical in five independent datasets. Junctions were assigned as canonical if their 5’ location was within 20 bases of the TRS-L and their 3’ location within 15 bases of a TRS-B, and otherwise assigned as non-canonical. Taiaroa, Kim, and Davidson are dRNAseq, while Finkel and Blanco-Melo are Illumina PolyA RNAseq. **B)** Illustration of computational approach to determine the consistency of 5’ and 3’ junction points across independent datasets. The percentage of non-canonical junctions at each genome position in each dataset was determined (X). The mean (μ) and standard deviation (α) of percentages at each position across the five datasets was calculated. For each position in each dataset, the Z-score was calculated as (X-μ)/α (i.e. the number of standard deviations away from the mean), and the percentages and Z-scores for each position in each dataset were plotted. **C, D)** Each point represents the percentage and Z-score of one position in one dataset. For each position in the SARS-CoV-2 genome, the percentage and Z-score of non-canonical junctions with a 5’ end (D) or a 3’ end (E) was determined as described above for five independent datasets: Taiaroa, Kim, Davidson, Finkel, and Blanco-Melo. Points with a percentage above 4% of non-canonical junctions and a Z-score above 1 were highlighted.

We sought to assess if non-canonical junctions are driven by dataset-specific artifacts. For example, it is possible that defective particles containing defective viral genomes (DVGs), often with large internal deletions that increase their replicative fitness, could accumulate during preparation of virion stocks (16) and generate the observed non¬canonical junctions during subsequent infections. If this was true, we would expect to observe non-canonical junctions specific to a single virion preparation and accumulation of these specific non-canonical junctions over the course of infection. To address this possibility, we identified dataset-specific junctions by identifying junctions that are differentially abundant across datasets (Figure 2B). This analysis revealed that the vast majority of 5’ and 3’ non-canonical junctions are low abundance and are similarly abundant across five independent SARS-CoV-2 sequencing studies (7–9, 12, 13) (Figure 2C, D). This suggests that most non-canonical junctions are not driven by dataset-specific artifacts.

However, we found that three 5’ junction sites in the Finkel et al. transcriptome and three 3’ junctions in the Davidson et al. transcriptome were overrepresented (Figure 2C, D). Incorporating additional replicates of these transcriptomes and assessing the percentage of non-canonical junctions at each site revealed that the Finkel et al.- specific 5’ junctions are highly elevated in both Finkel et al. replicates relative to other transcriptomes (Supplementary Figure 3A–C). In contrast, the three Davidson et al.- specific 3’ junctions are present in other datasets but at slightly lower frequency (Supplementary Figure 3D, E). The consistency of the Finkel et al.-specific junctions across replicates and exclusion from other samples raises the possibility that these junctions were generated from defective particles in the Finkel et al. virion preparation.

### The proportion of non-canonical junctions increases over time *in vitro*

Next, we assessed changes in junctions over time using the approach detailed in Figure 3A using transcriptomes from early (4 or 5 hours post infection) and late (24 hours post infection) timepoints from Finkel et al. (12) and Emanual et al. (14). This analysis revealed that there are large decreases in 5’ junctions originating at position 64, and in 3’ junctions ending in 28254 (proximal to the N TRS-B), with a smaller decrease in 3’ junctions ending at 26467 (proximal to the M TRS-B) (Figure 3B,C). Notably, the Finkel et al.-specific 5’ junction at 16971 increased in abundance over time. In contrast, there were no specific large increases in percentage of junctions at other positions (Figure 3B,C). Instead, there are small but widespread increases in non-canonical junction sites throughout the SARS-CoV-2 genome (Figure 3D, E). Quantification of the total percent changes of canonical and non-canonical junctions revealed a large decrease in canonical N junctions and increase in non-canonical junctions (Figure 3F, G). These results indicate that the abundance of non-canonical junctions increases over time *in vitro.* It is unclear if a similar pattern occurs during infection *in vivo.*

**Figure 3.**
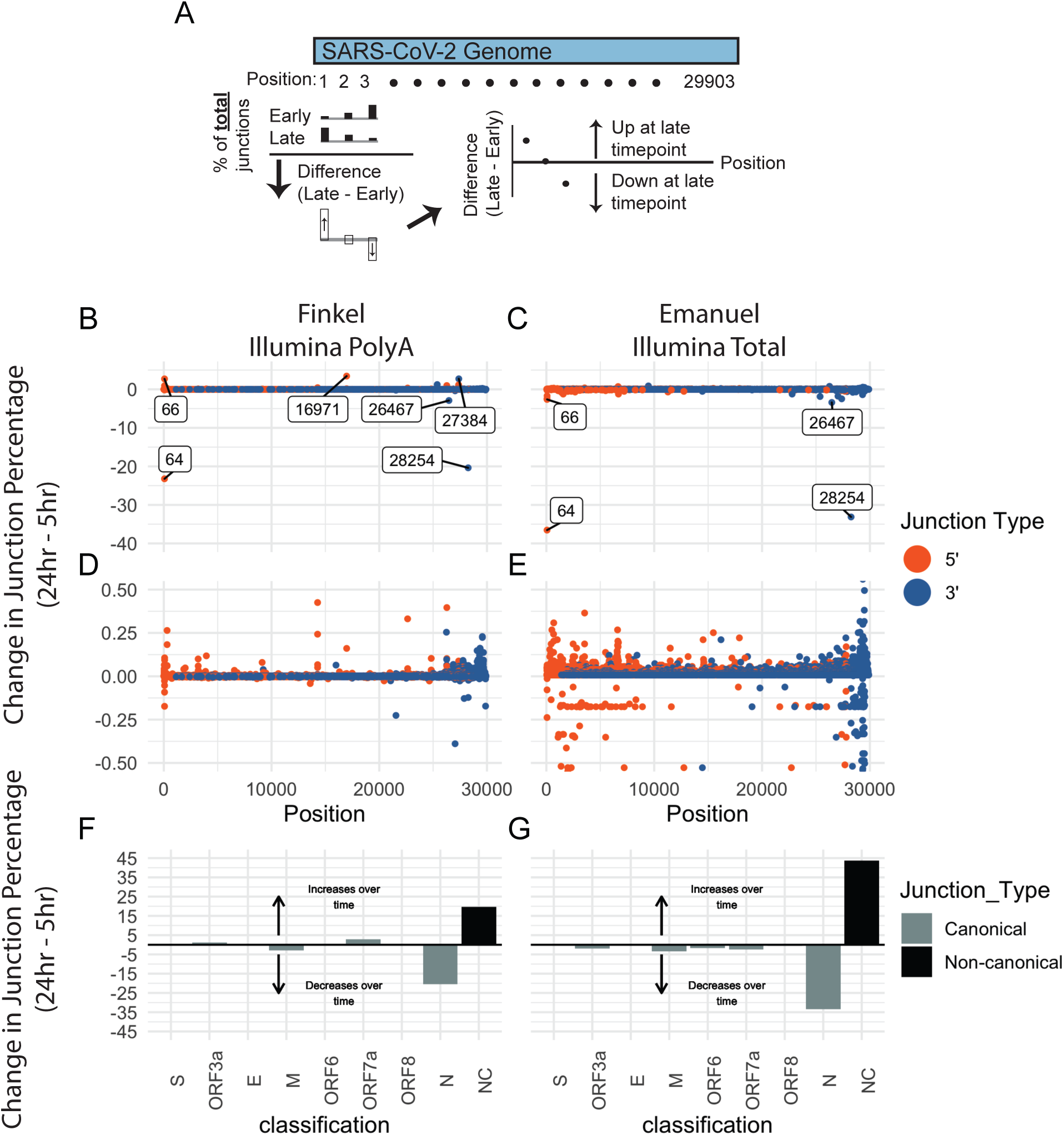
Non-canonical junctions accumulate over time *in vitro*. **A)** Illustration of computational approach used to determine change in junction percentages over time. For each position in the SARS-CoV-2 genome and separately for 5’ and 3’ junctions, the number of junctions at that position was calculated. Using these numbers, the percentage of 5’ or 3’ junctions falling at each position was calculated. The change in junction percentage at each position is defined as the difference between the position’s junction percentage at the late and early timepoints, and this junction change was plotted for each position. **B-E)** The change in junction percentage for 5’ (orange) and 3’ (blue) junctions over time in the Finkel (B, D) and Emanuel (C, E) datasets was determined as described above. Changes in the percentage of junctions Positions with at change greater than 2.5% are annotated with text on the plot. Panels D and E are zoomed in versions of B and C. **F, G)** Junctions were assigned into groups based on their 5’ and 3’ junction positions. If a junction had a 5’ end within 20 bases of the TRS-L and within 15 bases of a TRS-B, it was considered a canonical junction belonging to the ORF with a start closest to the TRS-B. Otherwise, it was considered non-canonical. The percentage of junctions falling into each category was calculated for early and late timepoints, and the difference between each category’s percentage in the late vs early timepoint was plotted.

### Non-canonical junctions are not associated with TRS-like homology

While homology between the TRS-L and TRS-B is thought to be critical for generation of canonical sgRNAs, the role of TRS-like homology in the generation of nc-sgRNAs is unclear. To address this question, we determined the 30 bases surrounding the 5’ and 3’ sites of each junction and identified the longest continuous subsequence in common between these sequence (Figure 4A). We found that junctions with a 3’ end landing within 15 nucleotides of a TRS-B overwhelmingly have junction homology associated with variants of the TRS core sequence (Figure 4B–D, **blue**). In contrast, junctions landing within ORFs and not proximal to a TRS-B do not have a prevalent homologous sequence (Figure 4B–D, **red**). The exception are intra-M junctions, whose homology is a consequence of the location of the ORF6 TRS within the M ORF.

**Figure 4.**
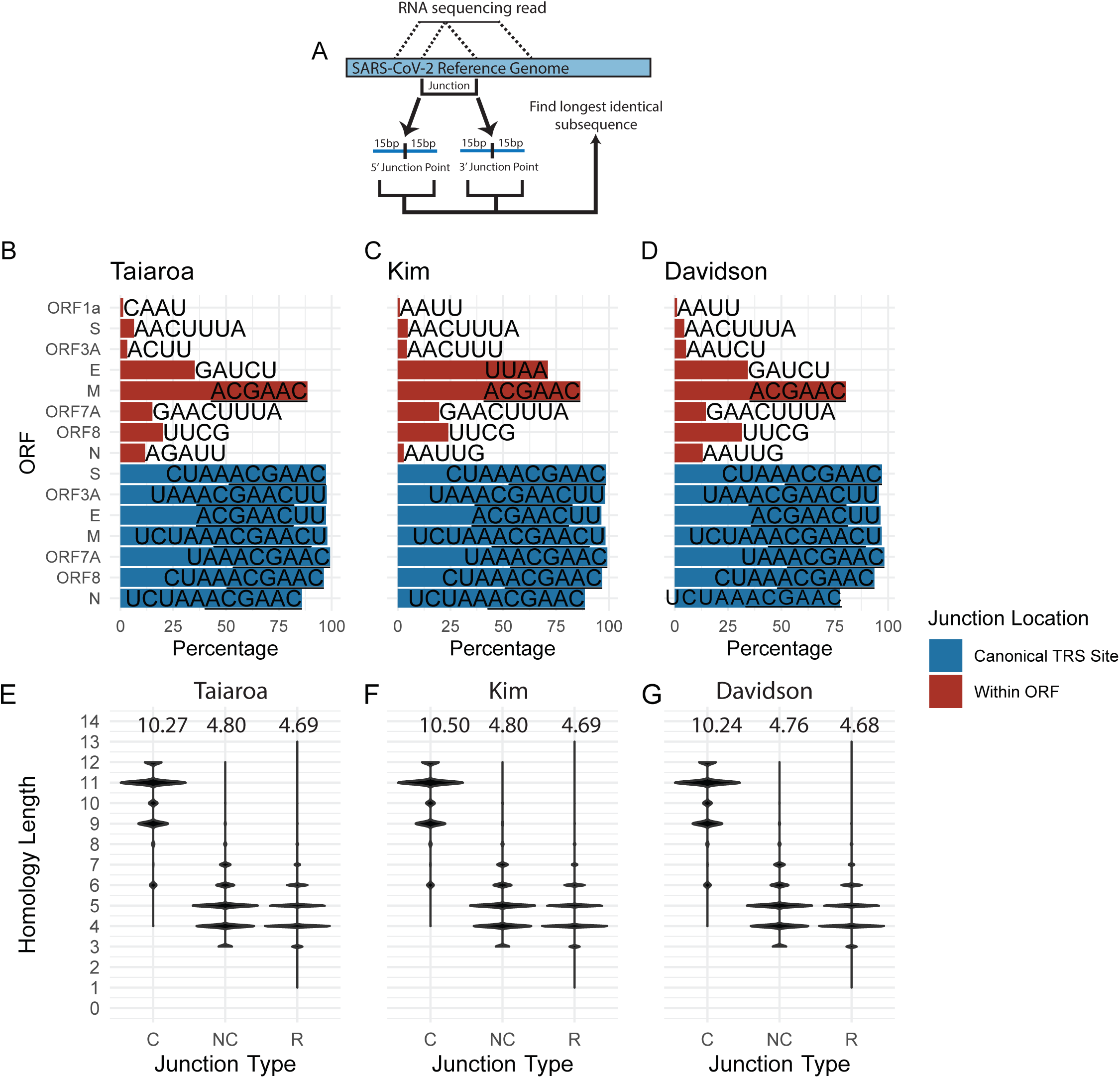
Non-canonical junctions are not associated with TRS-like homology. **A)** Illustration of computational approach used to assess homologous sequences between the 5’ and 3’ junction points. For each of the three dRNAseq datasets, the 30 bases flanking the 5’ and 3’ junction points were assessed, and the longest homologous sequence between these two 30 base regions was determined. **B-D)** Homologous sequences present in the 15 bases on either side of the 5’ and 3’ ends of each junction. Junctions are classified by the location of their 3’ end - if this is within 15 bases of the canonical TRS site or if it falls within a gene it is assigned accordingly. The only exception is ORF1a - junctions with a 5’ end originating in ORF1a are assigned to ORF1a. Labels represent the most common homologous sequence between the ends of each junction. The core TRS sequence ACGAAC is underlined. **E-G)** The length of the longest homologous sequence for canonical (C) and non¬canonical (NC) are plotted. Here, junctions were considered canonical if they have a 5’ end within 20 nucleotides of the TRS-L and a 3’ end within 15 nucleotides of a TRS-B. (R) represents random homology lengths. The points for (R) were calculated by first extracting all possible 30 base sequences (30mers) from the SARS-CoV-2 genome, and then assessing the length of the longest homologous sequence between 100,000 random pairs of 30mers that are separated by at least 1000 bases on the genome. Within each column, the relative width of each band represents the relative abundance of junctions with each homology length. The value at the top of each column is the mean homology length.

While there is not a single strong motif driving non-canonical, intra-ORF junctions, it is still possible that non-canonical junctions are associated with junction-specific homology between the 5’ and 3’ junction sites. To address this possibility, we tallied the length of homology between the 5’ and 3’ sites of canonical and non-canonical junctions. This analysis revealed that canonical junctions have markedly longer homology lengths (mean >10 bases) than non-canonical junctions (mean < 5 bases) (Figure 4E–G). Indeed, the homology lengths of non-canonical junctions are only slightly longer than homology lengths between 100,000 random pairs of SARS-CoV-2-derived 30 base sequences (Figure 4E–G). These data indicate that non-canonical junctions are not associated with TRS-like homology.

### sgRNAs containing only a 5’ region of ORF1a are produced in all studied *in vitro* and *in vivo* infection systems

Assessment of the three dRNAseq transcriptomes revealed evidence of sgRNAs that contain only the 5’ region of ORF1a (Supplementary Figure 3A). This is similar to an observation previously reported by Kim et al. (8). Inspection of read coverage of ORF1a in these transcriptomes confirmed that there is elevated 5’ ORF1a coverage with specific decreases in coverage concomitantly with sharp increases in junctions near genome positions 1600 and 6000 (Figure 5A–C).

**Figure 5.**
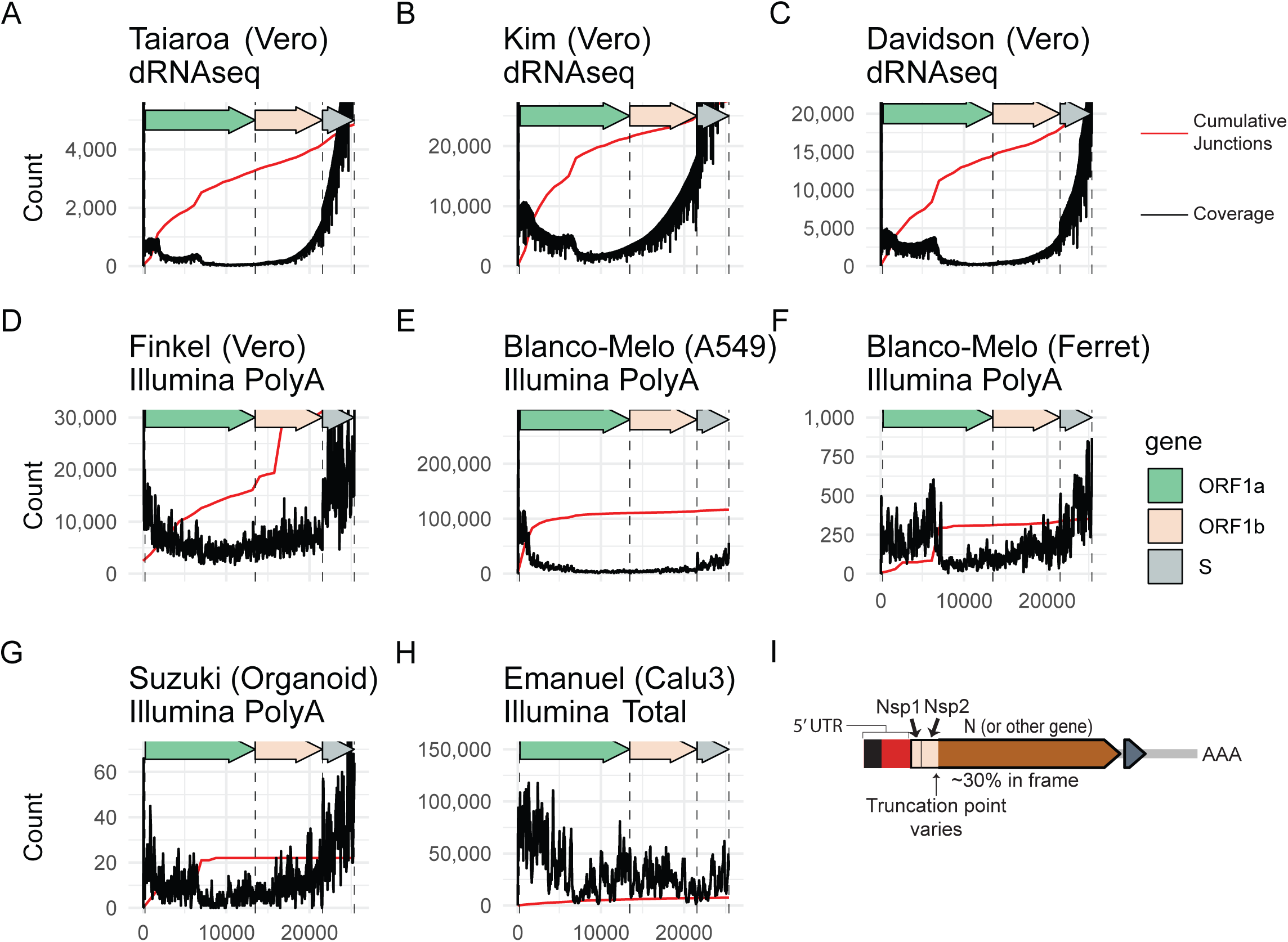
Relative abundance of subgenomic RNAs that contain only the 5’ end of ORF1a. **A-H)** Read coverage (black) and cumulative 5’ junctions (red) are plotted for eight independent datasets. The sequencing strategy and sample types are annotated. **I)** A schematic of a representative subgenomic RNA that consists of only the 5’ region of ORF1a. Pileups showing reads can be found in Supplementary Figure

While there is clearly higher coverage of the 5’ region of ORF1a in the three dRNAseq transcriptomes, it remained unclear if these 5’ ORF1a sgRNAs are a major percentage of total ORF1a in the cell, or if this finding was due to a systematic bias favoring shorter reads in nanopore sequencing. To address this question, we assessed ORF1a coverage in four independent Illumina polyA-enriched RNAseq transcriptomes. Analysis of these transcriptomes revealed a similar elevated 5’ ORF1a coverage and consistent drops in coverage around genome positions 1600 and 6000 (Figure 5D–G). Importantly, one of these transcriptomes is from bronchial organoids (Figure 5G) and one transcriptome is from the nasal washings of an infected ferret (Figure 5F), suggesting that 5’ ORF1a subgenomic RNAs can be produced *in vivo* and were not only due to artifacts of highly replicative conditions *in vitro.* Furthermore, analysis of a transcriptome generated from RNA that was not polyA-enriched revealed a consistent elevation in 5’ ORF1a coverage (Figure 5H), ruling out a systematic bias from polyA isolation. We found that there is similar elevated 5’ ORF1a coverage in the circulating human coronavirus CoV 229E (17) (Supplementary Figure 4), supporting the existence of 5’ ORF1a RNAs in other coronaviruses. All together, these data support the hypothesis that subgenomic RNAs containing just the 5’ region of ORF1a are produced by SARS-CoV-2.

### Junctions have the potential to generate variant open reading frames

To test the hypothesis that non-canonical junctions can result in variant open reading frames (ORFs) in nc-sgRNAs, we assessed the ORF content of each read in the three dRNAseq transcriptomes. Because dRNAseq can have an error rate of greater than 10% (18), we generated “coordinate-derived transcripts” by determining the start, end, and junction coordinates of each raw transcript by mapping to SARS-CoV-2 (NC_045512.2) and extracting the associated genomic sequence. While this approach is capable of overcoming sequencing errors and can resolve major RNA species, it is not capable of resolving small or structurally complex RNA rearrangements. Because the 5’ end of nanopore reads may be truncated due to technical factors during sequencing (19), we leveraged the assumption that functional sgRNAs should contain the 5’ leader sequence to identify reads generated from complete transcripts. While excluding reads that don’t contain the leader sequence may have reduced sensitivity, it prevents identification of variant ORFs as a consequence of truncated reads.

ORFs were predicted directly from the coordinate-derived transcripts, and ORFs of unexpected length were identified as “variant”. Assessment of the relative abundance of variant and canonical ORFs revealed evidence of variant ORFs of ORF1a/b, S, and M (Figure 6A).

**Figure 6.**
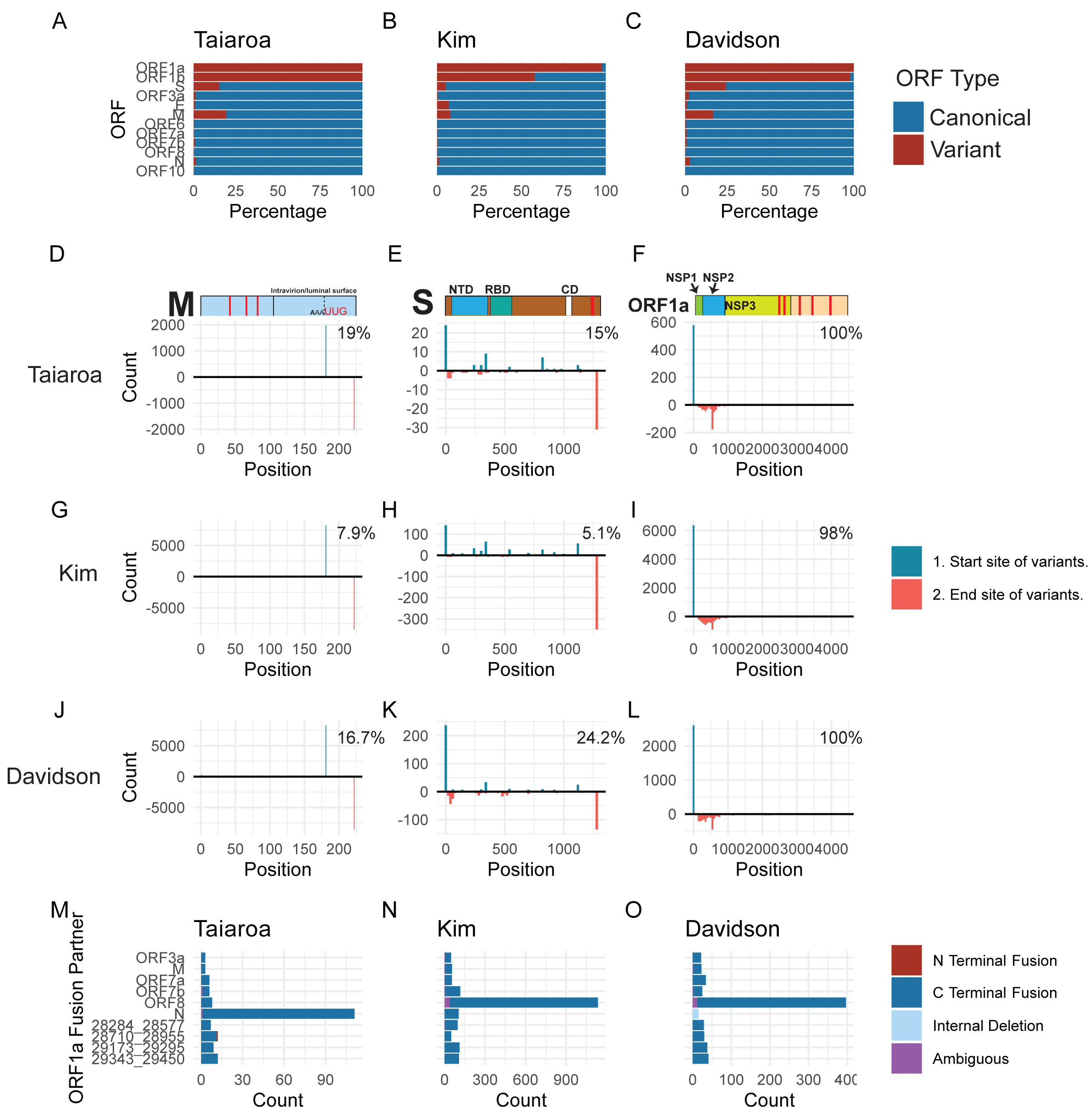
Junctions have the potential to generate variant open reading frames. **A-C)** ORFs were predicted directly from transcript-derived reads for the three dRNAseq datasets. Each ORF was aligned against the protein sequences of canonical SARS-CoV-2 genes using the DIAMOND aligner. Variant ORFs were defined as ORFs that were assigned to a canonical SARS-CoV-2 protein but had an unexpected start or stop position, while perfectly-aligning ORFs were considered canonical. The percentage of canonical and variant ORFs for each protein is plotted. **D-L**) Schematics of M, N, and ORFlab are displayed with the approximate location of predicted transmembrane domains labeled in red. A histogram of the start and end sites of variant M, N, and ORFlab ORFs are displayed. Start sites of each variant ORF are on top, and end sites are on the bottom of each panel. Percentages represent the percentage of each ORF that is variant. Histogram bin-sizes: M, 1; N, 10; ORFlab, 20. NTD: N-terminal domain. RBD: Receptor binding domain. CD: Connector domain. NSP: Nonstructural protein. **M-O)** The identity of ORF1a-fusion partners is plotted on the Y axis, with the count of such fusions on the X axis. The top 10 fusion partners for each sample are represented. Color indicates if the fusion partner is on the N and C terminus of ORF1a, if the terminus is ambiguous, or if the fusion is a “self” fusion between an upstream and downstream region of ORF1a. The percentage notes the percent of ORFs assigned to ORF1a that have a fusion.

To characterize the identified M, S, and ORF1a ORF variants, we determined the stop and start points of each ORF variant relative to their canonical counterparts (Figure 6D–L). This analysis revealed that all theoretical M ORF variants were predicted to start at a UUG start codon 42 amino acids before the end of the M gene and end at the canonical stop codon (Figure 6D, G, J). M ORF variants make up between 7.9% and 19% of M ORFs in the three dRNAseq datasets and are generated by the major junction point at the ORF6 TRS sequence within the M reading frame (Figure 1D–F). If translated, M ORF variants would lack the N-terminal transmembrane domains and contain regions predicted to be present on the inside of the virion or in the cytosol. While the efficacy of translation initiation from this start codon is unknown, it has an acceptable Kozak consensus site with an A at the -3 position.

S ORF variants make up between 5.1% and 24.2% of all S ORFs identified in these datasets (Figure 6A–C). The majority of S ORF variants are predicted from nc-sgRNAs with 3’ junctions landing within the S ORF (Supplementary Figure 2D–F), resulting in N-terminal truncations with theoretical AUG and non-AUG start codons (Figure 6E, H, K). If translated, these variants would lack the N-terminal signal sequence (20) and, especially considering the variations in ORF start sites, it is unclear if these ORFs encode functional isoforms.

Our identification of ORF1a variants is consistent with the observed patterns of 5’ ORF1a coverage (Figure 5). The peak end position of identified ORF1a variant ORFs is around amino acid position 500 (Figure 6F, I, L), consistent with the observed decrease in read coverage around genome position 1600. This position falls within the NSP2 amino acid sequence suggesting that, if translated, many ORF1a ORF variants would consist of NSP1 and N-terminal fragments of NSP2. Interestingly, we found that approximately 1/3 of ORF1a variants are in-frame fusions with downstream ORFs, encoding mostly N (Figure 6M–O).

## Discussion

While prior studies have identified nc-sgRNAs in SARS-CoV-2 (8, 11) and other coronaviruses (17), there has been no comprehensive analysis of SARS-CoV-2 nc-sgRNAs. We conducted an integrative analysis of eight independent SARS-CoV-2 transcriptomes to catalogue unexpected, non-canonical RNA species. We show that SARS-CoV-2 produces nc-sgRNAs, that these nc-sgRNAs increase in abundance over time, and that they are not associated with TRS-like homology. We highlight strong evidence supporting the existence of nc-sgRNAs containing only the 5’ region of ORF1a and show that the presence of nc-sgRNAs can alter the landscape of viral ORFs.

We found that canonical sgRNAs are consistently produced across SARS-CoV-2 transcriptomes, and identify eight major sgRNAs encoding S, ORF3a, E, M, ORF6, ORF7a/b, ORF8, and N. Each of these sgRNAs is supported by abundant 3’ junctions originating at a TRS-B just upstream the associated ORF. Similar to other reports, we do not find evidence of a major series of sgRNAs encoding ORF10 (7, 8, 11). We also do not find strong evidence of a major group of sgRNAs encoding ORF7B despite the presence of a TRS-core-like sequence (A**A**GAAC) proximal to the ORF7B start.

Interestingly, up to 33% of sgRNAs from infected cells *in vitro* are non-canonical. One potential source of nc-sgRNAs are DVGs introduced by defective particles during infection. These can accumulate during the propagation of virus stocks. While information on the passaging of viral stocks is only available for four transcriptomes, at least five of the six SARS-CoV-2 transcriptomes generated from cell lines were generated following a low MOI infection, which may reduce the impact of defective particles (21, 22). Indeed, while there is some variation in the absolute abundance of non-canonical junctions across transcriptomes that may reflect dataset-specific factors, the relative abundance of non-canonical junctions at most genomic positions is generally similar. One exception is that dataset-specific junctions are found in the Finkel et al. transcriptomes. In particular, there is a strong 5’ junction point at position 16971 that represents over 12% of non-canonical junctions in two Finkel et al. replicates but is largely absent in other transcriptomes. A hypothesis that could explain this observation is that there is a population of defective particles in the Finkel et al. virion preparation that contain an internal deletion. This DVG, which is smaller than the full genome, may have a replicative advantage during virus stock preparation (16). Indeed, the Finkel et al.-specific junction at 16971 displays a relatively large increase in abundance over time, in contrast to smaller and non-specific increases of non-canonical junctions at other positions.

The consistent presence of non-canonical junctions across the SARS-CoV-2 genome in independent transcriptomes suggests that the bulk of the observed nc-sgRNAs are not dataset-specific but are generalizable to SARS-CoV-2 biology. Furthermore, their nonspecific increase in abundance over time *in vitro* supports a model in which nc-sgRNAs are generated nonspecifically and at higher frequencies later in infection. We found that non-canonical junctions are not associated with TRS-like homology, raising the possibility that there is another mechanism behind their formation. For example, it is possible that factors such as RNA structure (23), accumulation of viral proteins, or actions of the viral RdRp and associated cofactors (24) could influence nc-sgRNA formation. Indeed, a recent study of the murine betacoronavirus MHV suggests that the 3’-to’5’ exoribonuclease Nsp14 influences recombination site selection of the viral RdRp (24). Additional studies are necessary to gain a mechanistic understanding of the factors that influence nc-sgRNA generation and to understand the phenotypic effects of nc-sgRNAs on SARS-CoV-2 pathogenesis. Notably, considering that defective viral genomes in negative-sense RNA viruses have been associated with antiviral immunity, dendritic cell maturation, and interferon production (5, 16, 25, 26) it will be important to assess if SARS-CoV-2 nc-sgRNAs are associated with immune activity.

Junctions in viral RNA change the landscape of viral ORFs. Because the ORF6 TRS-B is located 151 bases into the M ORF, sgRNAs created using this TRS-B contain a 3’ region of M. There is a UUG located 41 codons prior to the end of M that will now be situated prior to the ORF6 on these sgRNAs. While UUG is a relatively weak start codon, there is potential that this UUG could stimulate translation of a C-terminal M fragment or regulate the expression of the downstream ORF6 as an upstream ORF (27). Additional studies are needed to understand if this M ORF variant is translated and to understand if its UUG start codon influences ORF6 translation.

In addition to this M ORF variant, we found that some nc-sgRNAs contain S and ORF1a ORF variants. Many of the S ORF variants would lack the N-terminal signal sequence (28) and, especially considering the variation in the hypothetical start positions of these variants, it is unclear if they are translated. Notably, subgenomic RNAs encoding variant S ORFs have been reported for SARS-CoV-1 (29).

In contrast, the ORF1a ORF variants have a peak stop position around ORF1a amino acid position 500, which correlates well with an observed decrease in ORF1a 5’ coverage around genome position 1600. There is strong support for the existence of 5’ ORF1a-containing nc-sgRNAs across all eight transcriptomes studied here. Although there isn’t a strong peak of 5’ junctions at specific positions in the 5’ region of ORF1a, the consistent decreases in coverage and concomitant increases in 5’ junctions around positions 1600 and 6000 are notable. It is possible that there are biological factors at these positions that mediate these generalized inflection points but, unlike TRS-mediated homology, don’t concentrate junctions to a specific base. As posited by Kim et al. (8), the existence of subgenomic RNAs with the 5’ end of ORF1a raises the possibility that these nc-sgRNAs may influence the relative abundance of Nsps, making this observation an important target for future study.

## Conclusions

In conclusion, we show that SARS-CoV-2 generates nc-sgRNAs *in vitro* and *in vivo*, and that these nc-sgRNAs are largely consistent across independent transcriptomes. We show that nc-sgRNAs are not associated with TRS-like homology, suggesting there may be a homology-independent mechanism driving their formation. Finally, we show that non-canonical junctions have the potential to influence the landscape of viral ORFs, and that there is especially strong support for nc-sgRNAs that contain only a 5’ portion of ORF1a. Future studies are necessary to understand the mechanisms behind nc-sgRNA formation, and to understand the influence of nc-sgRNAs on SARS-CoV-2 pathogenesis.

## Materials and Methods

### Sequencing data

Details on all studied transcriptomes are listed in the table below:

| Source | Virus Isolate (MOI)<br>(Details on stock generation) | Infection<br>host | Timepoint | Accession or DOI | Technology |
| --- | --- | --- | --- | --- | --- |
| Taiaroa et al. (7) | Australia/VIC01/2020 (MOI: Unknown)<br>(Passage unknown) | Vero cells | Unknown | SRR11350376 | dRNAseq |
| Kim et al. (8) | Korea/KCDC02/2020 (MOI: 0.05)<br>(4<sup>th</sup> passage. No plaque isolation) | Vero cells | 24hpi | 10.17605/OSF.IO/8F6N9 | dRNAseq |
| Davidson et al. (9) | England/VE6-T/2020 (MOI: Unknown)<br>(2<sup>nd</sup> passage. No plaque isolation). | Vero cells | Unknown | 10.5281/zenodo.3722580 | dRNAseq |
| Finkel et al. (12) | GISAID Acc. No. EPI_ISL_406862 (MOI: 0.2)<br>(4<sup>th</sup> passage. No plaque isolation.) | Vero cells | 5hpi | SRR11713354 | Illumina<br>PolyA |
| Finkel et al. (12) | GISAID Acc. No. EPI_ISL_406862 (MOI: 0.2)<br>(4<sup>th</sup> passage. No plaque isolation.) | Vero cells | 24hpi | SRR11713362 | Illumina<br>PolyA |
| Blanco-Melo et al. (13) | USA/WA1/2020 (MOI: 0.2)<br>(Passage unknown) | A549-ACE2<br>cells | 24hpi | SRR11517741 | Illumina<br>PolyA |
| Blanco-Melo et al. (13) | USA/WA1/2020 (5*10<sup>4</sup> PFU/ferret)<br>(Passage unknown) | Ferret<br>(Nasal<br>washings) | 3dpi | SRR11517855-SRR11517858 | Illumina<br>PolyA |
| Suzuki et al. (15) | SARS-CoV-2/Hu/DP/Kng/19-020 (5*10<sup>4</sup> PFU/100 organoids)<br>(Passage unknown but **plaque isolated**) | Bronchial organoids | 5dpi | SRR11811022 | Illumina<br>PolyA |
| Emanuel et al. (14) | Frankfurt strain AY310120.1 **OR**<br>BetaCoV/Munich/BavPat1/2020\|EPI_ISL_406862<br>(MOI: 0.33)<br>(Passage unknown) | Calu3 | 4hpi | SRR11550047 | Illumina<br>Total RNA |
| Emanuel et al. (14) | Frankfurt strain AY310120.1 **OR**<br>BetaCoV/Munich/BavPat1/2020\|EPI_ISL_406862<br>(MOI: 0.33)<br>(Passage unknown) | Calu3 | 24hpi | SRR11550045 | Illumina<br>Total RNA |
| Viehweger<br>et al. (17) | CoV-229E (MOI: 3)<br>(Passage unknown) | Huh7<br>cells | 24hpi | ERR3460961 | dRNAseq |

Hpi: Hours post infection
Dpi: Days post infection

In the case of the Finkel et al. and Emanual et al. transcriptomes, the 24hpi samples were used in all analyses except for the investigation of junction changes over time.

### Determining junction coordinates

Reads in all transcriptomes were mapped against the SARS-CoV-2 reference genome (accession NC_045512.2) using minimap2 (30) and custom parameters detailed in minimap_sars2.sh and inspired by Kim et al. (8), and unmapped reads were discarded. The exceptions are the transcriptomes from Finkel et al. and Emanuel et al. - because their read lengths were very short (60 and 72 bases in length respectively), mapping was conducted with STAR (31) using settings detailed in star.sh.

The 5’ and 3’ junction positions of junction-spanning reads were determined with generate_synthetic_transcripts.py. This script reads a BAM file and the SARS-CoV-2 genome fasta and, by parsing the BAM alignment CIGAR and start position, determines the start, end, and junction coordinates of the alignment. Any minimap2-called deletion or intron of at least 1000 bases in length was considered a junction. Junction coordinates were then written to a tab-delimited output file for further analysis.

### Analysis and visualization of junctions

The 5’ and 3’ coordinates of each junction were processed in R (32), and figures generated with ggplot2 (33). Gene-maps were generated using the gggenes R package (34).

For the global distribution of reads in each transcriptome (Figure 1A–C, Supplementary Figure 1A–F), specific 5’-3’ junctions that occur at least twice are represented as arcs. Each inverted peak represents a histogram of 5’ or 3’ junctions at each genome position, with a bin size of 100 bases - this means that the magnitude of each peak represents the total number of 5’ or 3’ junctions within each 100 base span. Code: global_junctions.Rmd

The distribution of major 3’ junctions (Figure 1D–F) was generated by plotting a histogram of 3’ junction positions that occur after position 21000 and that have a 5’ end prior to position 100. Bin-size is 20 bases. Code: canonical_3prime_junctions.Rmd

The percentage of non-canonical junctions in each transcriptome (Figure 2A, Supplementary Figure 1G) was calculated by first assigning each individual 5’-3’ junction as canonical or non-canonical. A junction was considered to be canonical if it’s 5’ position was within 20 bases of the TRS-L core sequence ACGAAC (starting at position 69), and it’s 3’ junction was within 15 bases of a TRS-B (defined by the presence of the TRS core sequence). The percentage of total junctions that were assigned as non-canonical was then determined. Code: global_junctions.Rmd

### Identifying dataset-specific junctions (Figure 2)

The consistency of non-canonical junctions across transcriptomes (Figure 2C, D) was determined separately for 5’ and 3’ junctions based on one replicate each from Taiaroa et al., Kim et al., Davidson et al., Finkel et al., and Blanco-Melo et al. transcriptomes generated from cells infected with SARS-CoV-2 *in vitro.* For each SARS-CoV-2 genome position, the percentage of non-canonical junctions at that site was calculated for each transcriptome (X). The mean (μ) and standard deviation (σ) of percentages at each site across the five transcriptomes were then calculated. For each position in each transcriptome, a z-score was calculated by subtracting the observed percentage by the mean and dividing by the standard deviation ((X-μ)/σ). This process is visualized in Figure 2B. Code: dataset_specific_junctions.Rmd

### Change in junctions over time (Figure 3)

The change in junctions over time was determined based on datasets from Finkel et al. (5hpi and 24hpi) and Emanuel et al. (4hpi and 24hpi). The percentage of junctions that originate at each genome position was determined for each timepoint, and the absolute difference between the two values was calculated. Notably, this analysis (Figure 3B, C) focused on 5’ and 3’ counts at each position separately, such that a percentage of 5’ junctions and a percentage of 3’ junctions at each position was determined. To determine the aggregated change in each <u>class</u> of junctions (Figure 3F,G), each 5’-3’ junction was assigned to a class based on the location of its 5’ and 3’ position. If a junction’s 5’ position was located within 20 bases of the TRS-L, and it’s 3’ position was located within 15 bases of a TRS-B, it was assigned to the ORF closest to the TRS-B. Otherwise, the junction was assigned as non-canonical. The percentage of total junctions in each class at the early timepoint was subtracted from the class percentages at the late timepoint for each dataset. Code: junctions_over_time.Rmd

### Assessing homology near 5’ and 3’ positions of each junction (Figure 4)

The identity and length of homologous sequences flanking the 5’ and 3’ sites of each junction were determined by identifying the 30 bases flanking each junction site. The longest identical subsequence between these two 30 base sequences was then collected. For Figure 4B–D, the most common homology sequence was plotted for each group of junctions. Here, a junction was considered canonical if it has a 3’ end within 15 bases of a canonical TRS-B or was assigned as “Within ORF” if it was otherwise within an ORF. The exception were junctions with a 5’ end within ORF1a - these were assigned to ORF1a. The total distribution of homology lengths was also determined for canonical and non-canonical junctions (Figure 4E–G). To determine the random distribution of homology lengths, all 30 base sequences (30mers) present in the SARS-CoV-2 genome were determined. The longest identical subsequence between 100,000 random pairs of 30mers that are derived from at least 1000 bases apart on the genome was determined. Code: TRS_plotting.Rmd

### Assessment of excess 5’ ORF1a genome coverage (Figure 5)

Coverage at each position was assessed from SARS-CoV-2-mapped BAMs using the depth subcommand of samtools (35). (Command: samtools depth -aa -d0 file.bam > file.cov). Coverage and cumulative number of 5’ junctions (excluding junctions occurring before position 100) were plotted. Code: ORF1a_coverage.Rmd

### Analysis of ORFs on dRNAseq reads (Figure 6)

generate_synthetic_transcripts.py was used to generate “coordinate-derived transcripts” from the SARS-CoV-2 mapped BAMs from the Taiaroa et al., Kim et al., and Davidson et al. dRNAseq transcriptomes. This script reads a BAM file and the SARS-CoV-2 genome fasta and, by parsing the BAM alignment CIGAR and start position, determines the start, end, and junction coordinates of the alignment. Coordinate-derived transcripts were created by using these coordinates to retrieve the associated sequence from the SARS-CoV-2 genome. Any minimap2-called deletion or intron of at least 1000 bases in length was considered a junction. All coordinate-derived transcripts were then aligned against a minimal leader sequence fragment (5’ - CTTCCCAGGTAACAAACCAACCAACTTTCGATCTCTTGTAGATCTGTTCTC - 3’) using blastn (36). This minimal sequence was used instead of the full leader sequence because inspection of 5’ coverage in the three long-read datasets revealed a significant drop in coverage at the end of this sequence. Transcripts with an alignment of at least 45 nucleotides in length and a percent identity of at least 85%, and that contain the start of this sequence within the first 50 nucleotides of the transcript, were kept. Transcripts not containing this leader sequence were discarded.

ORFs were called directly from each transcript using Prodigal (37), using parameters listed in prodigal.sh. ORFs were allowed to initiate from any nUG start codon, were not required to be the most 5’ ORF on each transcript and were only reported if they are at least 30 codons in length. Each ORF was translated into amino acid sequence using prodigal_to_orf_direct.py. If multiple Prodigal-predicted ORFs contained an overlapping amino acid sequence, the longest ORF was output. Each ORF was output in fasta format and labeled with transcript of origin and coordinates of the ORF on the transcript.

The amino acid sequence of each ORF was then mapped against canonical SARS-CoV-2 proteins (from accession NC_045512.2) as well as Prodigal-predicted unannotated ORFs from the SARS-CoV-2 genome using DIAMOND (38). DIAMOND parameters are described in diamond.sh and include an E-value threshold of 10. During downstream analysis, alignments were considered valid if the aligned portions maintained at least 95% amino acid identity to the reference protein.

Various statistics of each ORF, including alignment information, the start position of the ORF on the transcript, and fusion information, were generated from the DIAMOND alignment file using parse_orf_assignments.py. An ORF was considered variant if it was not the same length as the canonical protein, with a notable exception. Because there are alternative start codons upstream of some annotated genes, and Prodigal would call ORFs using these start codons if they are present on a transcript, ORFs were allowed to have up to 20 amino acids of additional N-terminal amino acids while still being classified as canonical if they end at the expected position. If an ORF had matches against multiple SARS-CoV-2 proteins, it was given a primary assignment of the protein with the longest alignment length.

## Code availability

The reproducible computational pipeline used to generate all results is publicly available at https://qithub.com/inoms/virORFdirect.

## List of abbreviations

CoV: Coronavirus
dRNAseq: Direct RNA sequencing
E: Envelope protein
M: Matrix protein
N: Nucleocapsid protein
Nsp: Nonstructural protein
ORF: Open reading frame
S: Spike protein
SARS-CoV-2: Systemic acute respiratory syndrome Coronavirus 2
TRS: Transcriptional regulatory sequence

## Declarations

### Ethics approval and consent to participate

Not applicable

### Consent for publication

Not applicable

### Availability of data and materials

Code is available in the virORF_direct repository, https://qithub.com/inoms/virORFdirect.

### Competing interests

M.M. receives research support from Bayer, Ono, and Janssen, has patents licensed to Bayer and Labcorp, and is a consultant for OrigiMed. J.A.D. received research support from Constellation Pharmaceuticals and is a consultant for EMD Serono, Inc. and Merck & Co. Inc.

## Funding

This work was supported in part by the US Public Health Service grants R35CA232128 and P01CA203655 to J.A.D.

## Authors’ Contributions

J.N. analyzed the data; J.N., M.M., J.A.D. designed the study and wrote the manuscript.

## Acknowledgements

The authors thank Taiaroa et al., Kim et al., Davidson et al., Blanco-Melo et al., Suzuki et al., Finkel et al., and Emanuel et al. for collecting the RNA sequencing data and making these data publicly available. Portions of this research were conducted by using the O2 High Performance Compute Cluster, supported by the Research Computing Group at Harvard Medical School.

---

**Supplementary Figure 1.**
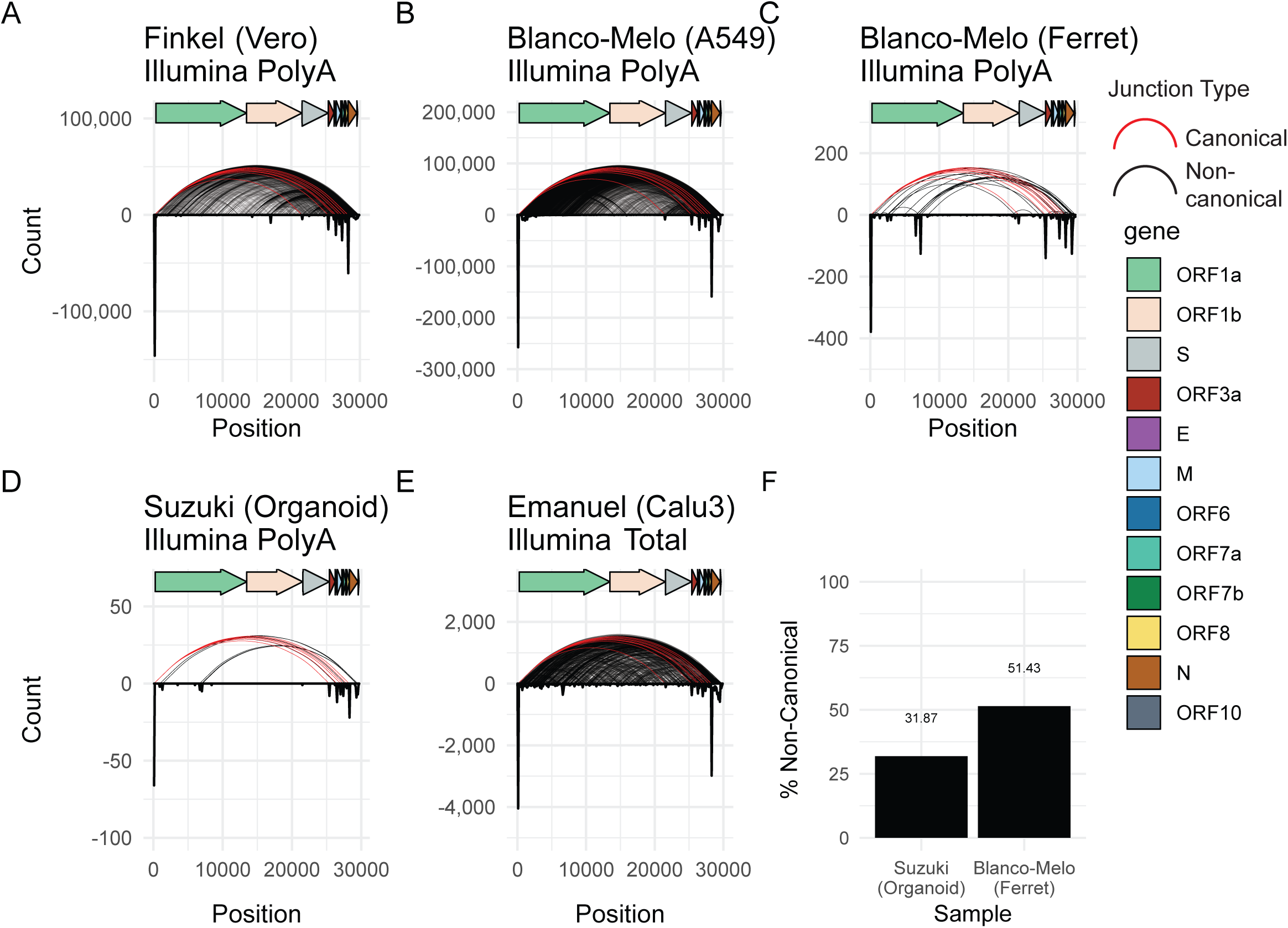
There is evidence of non-canonical junctions in diverse datasets. **A-E)** For each location on the viral genome, a histogram of 5’ and 3’ junctions at that position was calculated and plotted as an inverse peak. The histogram bin size is 100 bases, meaning each inverse peak represents the cumulative count of 5’ or 3’ junctions occurring within that span. Curved lines represent the 5’ and 3’ locations of junctions that occur at least twice. Red curves represent canonical junctions, black curves represent non-canonical junctions. **F)** The percentage of junctions that are canonical in the Suzuki (Bronchial organoids) and Blanco-Melo (Ferret) datasets was determined. Junctions were assigned as canonical if their 5’ location was within 20 bases of the TRS-L and their 3’ location within 15 bases of a TRS-B, and otherwise assigned as non¬canonical.

**Supplementary Figure 2.**
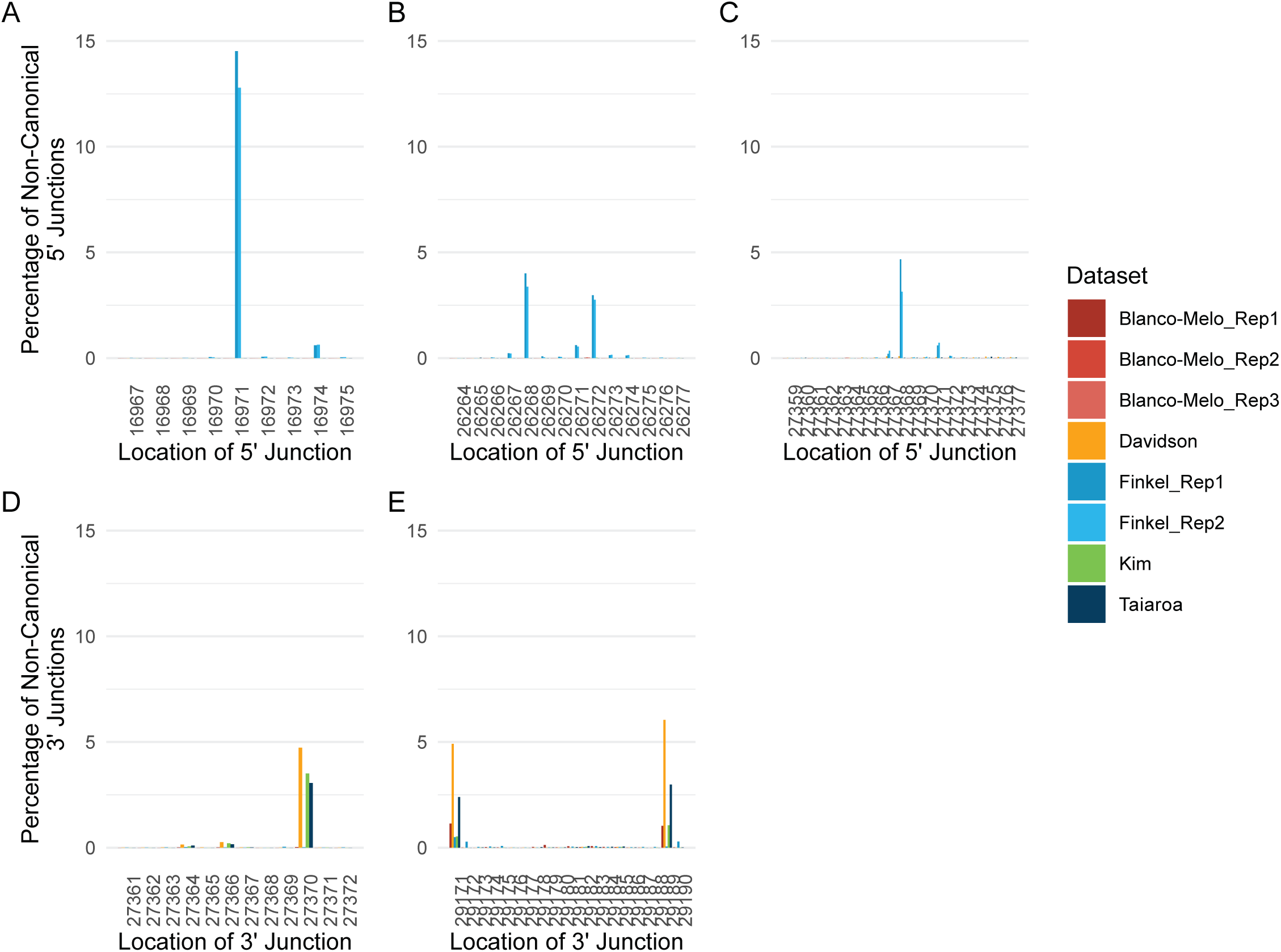
Dataset-specific junctions. **A-E)** The percentage of non-canonical junctions at each SARS-CoV-2 genome position is plotted. Plots A-C show the percentage of 5’ junctions at each position, while D-E show the percentage of 3’ junctions at each position. The plotted samples include three replicates from Blanco-Melo et al., Davidson et al, two replicates from Finkel et al, and the Kim and Taiaroa et al datasets.

**Supplementary Figure 3.**
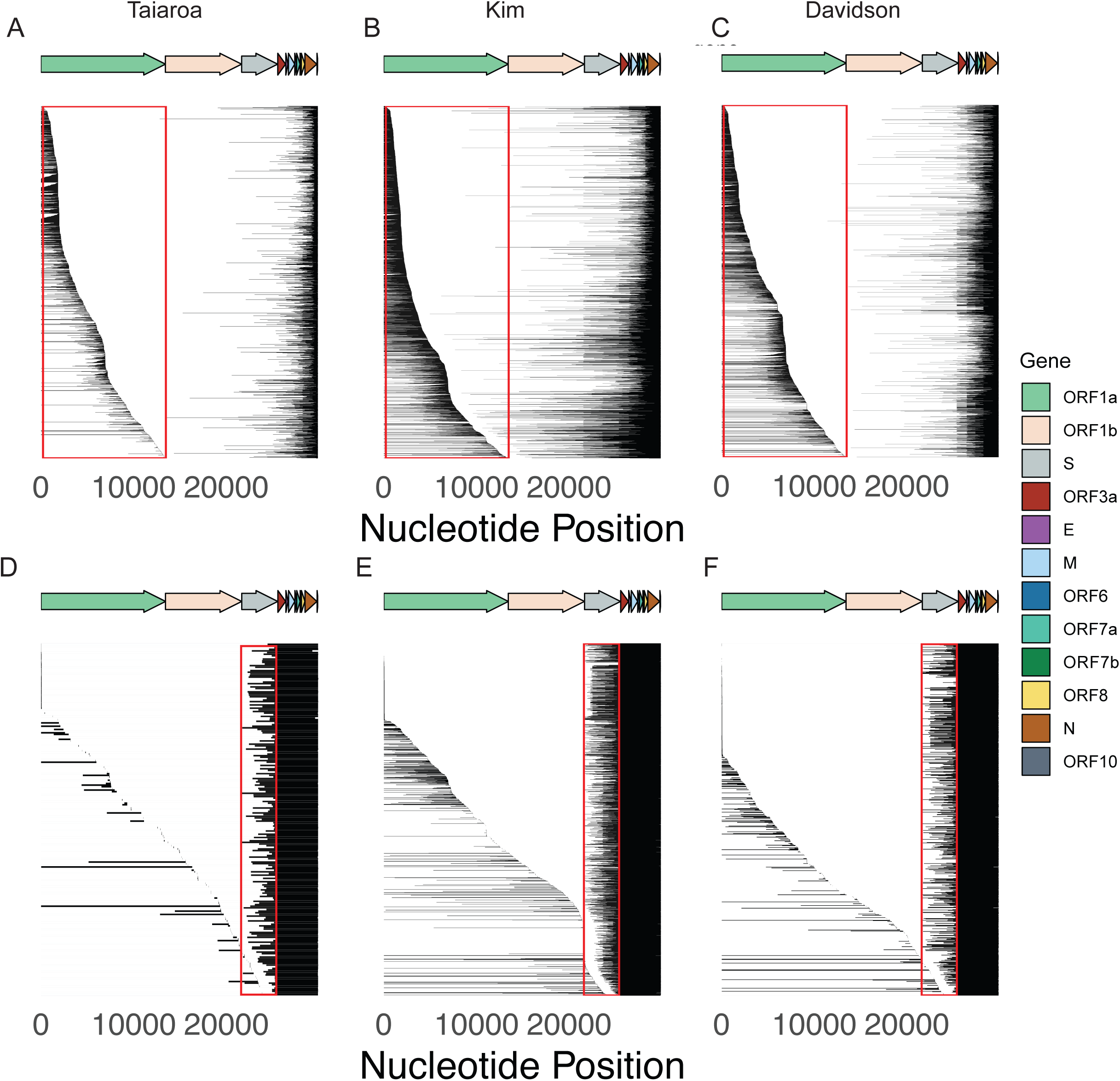
Subgenomic RNAs containing non-canonical junctions are present in three independent dRNAseq datasets. **A-F)** Subgenomic RNAs with 5’ junctions occurring in ORF1a (A-C) or 3’ junctions occurring in S (D-F) are plotted for the three dRNAseq datasets. Each row is a read, and regions in black indicate the read contains that sequence.

**Supplementary Figure 4.**
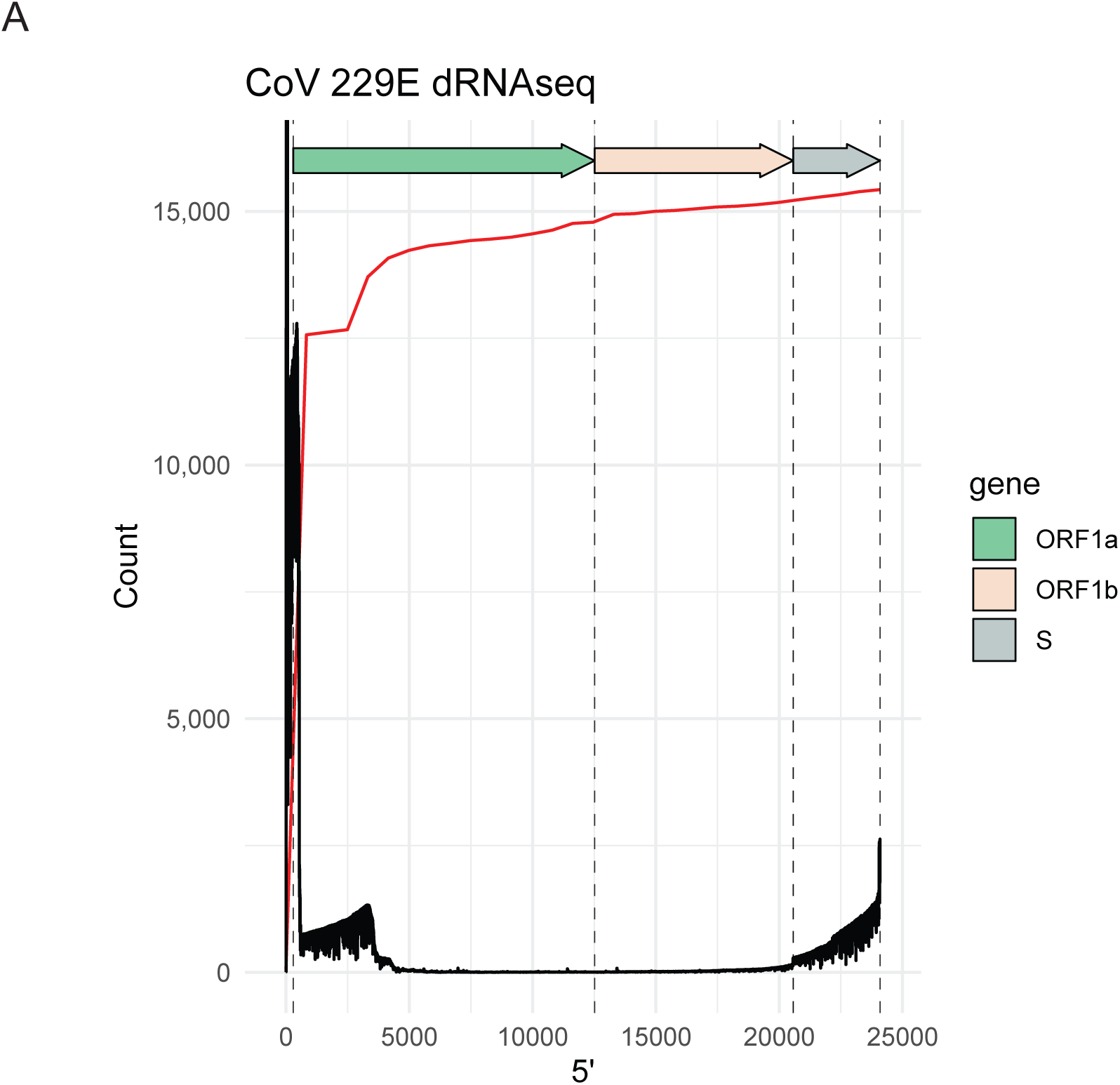
There is elevated ORF1a coverage in CoV 229E. **A)** Read coverage (black) and cumulative 5’ junctions (red) are plotted for dRNAseq data from CoV 229E-infected cells.

## References

1. Wu F, Zhao S, Yu B, Chen Y-M, Wang W, Song Z-G, et al. A new coronavirus associated with human respiratory disease in China. Nature. 2020;579(7798):265–9.

2. Plant EP, Dinman JD. The role of programmed-1 ribosomal frameshifting in coronavirus propagation. Frontiers in bioscience: a journal and virtual library. 2008;13:4873.

3. Narayanan K, Huang C, Makino S. SARS coronavirus accessory proteins. Virus research. 2008;133(1):113–21.

4. de Haan CA, Masters PS, Shen X, Weiss S, Rottier PJ. The group-specific murine coronavirus genes are not essential, but their deletion, by reverse genetics, is attenuating in the natural host. Virology. 2002;296(1):177–89.

5. Yount B, Roberts RS, Sims AC, Deming D, Frieman MB, Sparks J, et al. Severe acute respiratory syndrome coronavirus group-specific open reading frames encode nonessential functions for replication in cell cultures and mice. Journal of virology. 2005;79(23):14909–22.

6. Sawicki SG, Sawicki DL, Siddell SG. A contemporary view of coronavirus transcription. Journal of virology. 2007;81(1):20–9.

7. Taiaroa G, Rawlinson D, Featherstone L, Pitt M, Caly L, Druce J, et al. Direct RNA sequencing and early evolution of SARS-CoV-2. bioRxiv. 2020:2020.03.05.976167.

8. Kim D, Lee J-Y, Yang J-S, Kim JW, Kim VN, Chang H. The architecture of SARS-CoV-2 transcriptome. Cell. 2020.

9. Davidson AD, Williamson MK, Lewis S, Shoemark D, Carroll MW, Heesom KJ, et al. Characterisation of the transcriptome and proteome of SARS-CoV-2 reveals a cell passage induced in-frame deletion of the furin-like cleavage site from the spike glycoprotein. Genome Medicine. 2020;12(1):1–15.

10. Kim D, Lee J-Y, Yang J-S, Kim JW, Kim VN, Chang H. The architecture of SARS-CoV-2 transcriptome. bioRxiv. 2020:2020.03.12.988865.

11. Davidson AD, Williamson MK, Lewis S, Shoemark D, Carroll MW, Heesom K, et al. Characterisation of the transcriptome and proteome of SARS-CoV-2 using direct RNA sequencing and tandem mass spectrometry reveals evidence for a cell passage induced in-frame deletion in the spike glycoprotein that removes the furin-like cleavage site. bioRxiv. 2020:2020.03.22.002204.

12. Finkel Y, Mizrahi O, Nachshon A, Weingarten-Gabbay S, Yahalom-Ronen Y, Tamir H, et al. The coding capacity of SARS-CoV-2. bioRxiv. 2020:2020.05.07.082909.

13. Blanco-Melo D, Nilsson-Payant BE, Liu W-C, Uhl S, Hoagland D, Møller R, et al. Imbalanced host response to SARS-CoV-2 drives development of COVID-19. Cell. 2020.

14. Emanuel W, Kirstin M, Vedran F, Asija D, Theresa GL, Roberto A, et al. Bulk and single-cell gene expression profiling of SARS-CoV-2 infected human cell lines identifies molecular targets for therapeutic intervention. bioRxiv. 2020:2020.05.05.079194.

15. Suzuki T, Itoh Y, Sakai Y, Saito A, Okuzaki D, Motooka D, et al. Generation of human bronchial organoids for SARS-CoV-2 research. bioRxiv. 2020:2020.05.25.115600.

16. Genoyer E, López CB. The impact of defective viruses on infection and immunity. Annual review of virology. 2019;6:547–66.

17. Viehweger A, Krautwurst S, Lamkiewicz K, Madhugiri R, Ziebuhr J, Hölzer M, et al. Direct RNA nanopore sequencing of full-length coronavirus genomes provides novel insights into structural variants and enables modification analysis. Genome research. 2019;29(9):1545–54.

18. Rang FJ, Kloosterman WP, de Ridder J. From squiggle to basepair: computational approaches for improving nanopore sequencing read accuracy. Genome biology. 2018;19(1):90.

19. Workman RE, Tang AD, Tang PS, Jain M, Tyson JR, Razaghi R, et al. Nanopore native RNA sequencing of a human poly (A) transcriptome. Nature Methods. 2019;16(12):1297–305.

20. Bosch BJ, Martina BE, Van Der Zee R, Lepault J, Haijema BJ, Versluis C, et al. Severe acute respiratory syndrome coronavirus (SARS-CoV) infection inhibition using spike protein heptad repeat-derived peptides. Proceedings of the National Academy of Sciences. 2004;101(22):8455–60.

21. Thompson KAS, Yin J. Population dynamics of an RNA virus and its defective interfering particles in passage cultures. Virology journal. 2010;7(1):257.

22. Brooke CB. Biological activities ofnoninfectious' influenza A virus particles. Future virology. 2014;9(1):41–51.

23. Lan TCT, Allan MF, Malsick LE, Khandwala S, Nyeo SSY, Bathe M, et al. Structure of the full SARS-CoV-2 RNA genome in infected cells. bioRxiv. 2020:2020.06.29.178343.

24. Gribble J, Pruijssers AJ, Agostini ML, Anderson-Daniels J, Chappell JD, Lu X, et al. The coronavirus proofreading exoribonuclease mediates extensive viral recombination. bioRxiv. 2020:2020.04.23.057786.

25. Sun Y, Jain D, Koziol-White CJ, Genoyer E, Gilbert M, Tapia K, et al. Immunostimulatory defective viral genomes from respiratory syncytial virus promote a strong innate antiviral response during infection in mice and humans. PLoS Pathog. 2015;11(9):e1005122.

26. Strahle L, Garcin D, Kolakofsky D. Sendai virus defective-interfering genomes and the activation of interferon-beta. Virology. 2006;351(1):101–11.

27. Barbosa C, Peixeiro I, Romão L. Gene expression regulation by upstream open reading frames and human disease. PLoS Genet. 2013;9(8):e1003529.

28. Wrapp D, Wang N, Corbett KS, Goldsmith JA, Hsieh C-L, Abiona O, et al. Cryo-EM structure of the 2019-nCoV spike in the prefusion conformation. Science. 2020;367(6483):1260–3.

29. Hussain S, Chen Y, Yang Y, Xu J, Peng Y, Wu Y, et al. Identification of novel subgenomic RNAs and noncanonical transcription initiation signals of severe acute respiratory syndrome coronavirus. Journal of virology. 2005;79(9):5288–95.

30. Li H. Minimap 2: pairwise alignment for nucleotide sequences. Bioinformatics. 2018;34(18):3094–100.

31. Dobin A, Davis CA, Schlesinger F, Drenkow J, Zaleski C, Jha S, et al. STAR: ultrafast universal RNA-seq aligner. Bioinformatics. 2013;29(1):15–21.

32. Team RC. R: A language and environment for statistical computing. 2013.

33. Wickham H. ggplot2. Wiley Interdisciplinary Reviews: Computational Statistics. 2011;3(2):180–5.

34. Wilkins D. gggenes: draw gene arrow maps in ‘ggplot2’. R package version 0.4.0. 2019.

35. Li H, Handsaker B, Wysoker A, Fennell T, Ruan J, Homer N, et al. The sequence alignment/map format and SAMtools. Bioinformatics. 2009;25(16):2078–9.

36. Camacho C, Coulouris G, Avagyan V, Ma N, Papadopoulos J, Bealer K, et al. BLAST+: architecture and applications. BMC bioinformatics. 2009;10(1):421.

37. Hyatt D, Chen G-L, LoCascio PF, Land ML, Larimer FW, Hauser LJ. Prodigal: prokaryotic gene recognition and translation initiation site identification. BMC bioinformatics. 2010;11(1):119.

38. Buchfink B, Xie C, Huson DH. Fast and sensitive protein alignment using DIAMOND. Nature methods. 2015;12(1):59.

